# Do pyrethroid insecticides influence sex ratios in agile frogs (*Rana dalmatina*)?

**DOI:** 10.1101/2024.09.21.614238

**Authors:** Emese Balogh, Szabolcs Hócza, Nikolett Ujhegyi, Andrea Kásler, Dóra Holly, Dávid Herczeg, János Ujszegi, Zoltán Gál, Orsolya I. Hoffmann, Veronika Bókony, Zsanett Mikó

## Abstract

Environmental pollutants have the potential to alter sex ratios in wildlife through sex-biased mortality. Furthermore, endocrine disruptors may cause sex reversal during early ontogeny in ectothermic vertebrates, resulting in a phenotypic sex that is not concordant with the genotypic sex encoded by the sex chromosomes. Despite the wide-ranging implications of these sex-ratio biasing effects, they are rarely studied in ecotoxicology, especially in a way that allows for disentangling the two mechanisms. We investigated these effects of two synthetic pyrethroids, deltamethrin and etofenprox, that are commonly used insecticides and have been linked to adverse effects on fish and amphibian biodiversity. We assessed the effects of environmentally relevant concentrations of these two pyrethroids on phenotypic sex ratio, sex-dependent mortality, and sex reversal in agile frogs (*Rana dalmatina*). Tadpoles from field-collected eggs were reared in mesocosm until metamorphosis by adding 0.03 or 0.3 μg/L of deltamethrin or etofenprox three times to the water. We observed no effect in three of the four treatment groups. However, in the lower-concentration etofenprox treatment, phenotypic sex ratio was male-biased two months post-metamorphosis, and genotypic sexing revealed that this was due to female-biased mortality during metamorphosis and not to sex reversal. Although the estimation certainty of these effects was somewhat limited, they highlight that not all sex-ratio distorting effects are caused by sex reversal. Therefore, ecotoxicological studies aiming to understand the endocrine distruptor effects of environmental contaminants should strive to separate the effects on sex determination and sex-dependent mortality.

## INTRODUCTION

Sex ratio is a crucial apical (organismal) endpoint in ecotoxicology. The importance of this trait lies in its ultimate impact on population viability and adaptability, which is critical for biodiversity conservation. A skewed sex ratio can compromise a population’s ability to reproduce successfully and maintain genetic diversity, leading to decreased resilience and increased risk of extinction (Mitchell and Janzen, 2010). From a proximate perspective, changes in the sex ratio in ecotoxicological experiments can indicate endocrine disruption. Endocrine-disrupting chemicals (EDCs) interfere with the hormonal system, affecting somatic and sexual development, and posing a significant threat to wildlife across various species (Bergman et al., 2013; Orton and Tyler, 2015). While hundreds of environmental pollutants are known or suspected to be EDCs, *in vivo* tests of apical endpoints and especially of sex ratio have been done only on a fraction of these substances on a fraction of extant species (Bergman et al., 2013; Orton and Tyler, 2015).

Several mechanisms can lead to skewed sex ratios. First, one sex may be more sensitive to environmental stressors than the other, resulting in different mortality in males and females (Afonso et al., 2003). For instance, in a polluted environment, the mortality rate of lake char (*Salvelinus umbla)* embryos is higher in females than in males (Nusbaumer et al., 2021). Second, in numerous species, the development of offspring sex (i.e. sex determination) is susceptible to environmental influences. Several EDCs have been shown to affect offspring sex ratio in species with environmental sex determination, whereby the external environment primarily determines the sex of the developing individual (Mizoguchi and Valenzuela, 2016). In species with genotypic sex determination, exposure to environmental stressors such as chemical pollutants early in development (typically in the larval or embryonic stage of fish, amphibians, and reptiles) can cause sex reversal, meaning that the gonads that develop in the individual do not correspond to the genotypic sex determined by the sex chromosomes or other elements of the genome (Flament, 2016; Holleley et al., 2016; Nemesházi and Bókony, 2022; Ospina-Álvarez and Piferrer, 2008). These two mechanisms, i.e., sex-biased mortality *versus* altered sex determination, have rarely been separated and tested for in ecotoxicological experiments; instead, conclusions were often based solely on phenotypic sex ratios (Nemesházi and Bókony, 2022). A main reason for this shortcoming is that identification of genotypic sex is required both for demonstrating sex reversal and for analysing sex differences in mortality among young offspring where phenotypic sex is not yet detectable. Because of the evolutionary lability of sex determination across ectothermic vertebrates, genotypic sexing methods often need to be developed on a species-by-species basis; therefore, such methods are available only for a tiny proportion of extant taxa so far (Nemesházi and Bókony, 2022). Nevertheless, it is important to identify the underlying mechanisms of sex-ratio distorting endocrine disruptors, because the mechanisms influence the ultimate consequences on the long-term sustainability and evolutionary adaptability of wildlife populations (Geffroy and Wedekind, 2020). While both processes (i.e. sex-biased mortality and disrupted sex determination) skew the sex ratio, additionally sex-biased mortality may also put an extra burden on population viability by selectively killing the members of one sex. In contrast, sex reversal may impact individual survival and reproductive success (Holleley et al., 2015; Mikó et al., 2021; Nemesházi et al., 2021; Wild et al., 2022) and theoretical models indicate that these changes could have significant effects at the level of populations, including the evolution of sex-determination systems and mating preferences, and either reduced or heightened extinction risks depending on circumstances (Bókony et al., 2017; Grossen et al., 2011; Nemesházi et al., 2021; Schwanz et al., 2020). Therefore, it is crucial to differentiate between sex-specific mortality rates and sex reversal in EDC studies, because identifying the correct mechanism ensures that mitigation efforts target the appropriate causes of sex-ratio distortion, whether through reducing exposure to harmful pollutants or through interventions aimed at bolstering the survival of the more vulnerable sex (Geffroy and Wedekind, 2020).

Pyrethroids, which are commonly used as insecticides, are also among the endocrine-disrupting substances. They are effective substitutes for organochlorines and organophosphates (Gajendiran and Abraham, 2018). In comparison to the latter, pyrethroids have lower persistence in the environment and are relatively safer for birds and humans (Eni et al., 2019). However, they exhibit high toxicity to aquatic invertebrates (Palmquist et al., 2012) and fish (Haya, 1989). Pyrethroid insecticides are used in a wide range of applications, including agricultural, household, and veterinary use. They are also the primary active ingredients in chemical mosquito control. As a result, pyrethroids can enter surface waters, with measured concentrations typically ranging from 0.01 to 1 μg/L (Tang et al., 2018). One of the older pyrethroids, deltamethrin, is among the most widely used insecticides globally (Huang et al., 2014). Previous studies have demonstrated that sublethal concentrations of deltamethrin can be toxic to both fish and amphibians (Huang et al., 2014). Furthermore, it can influence their behaviour, induce developmental abnormalities, and even cell death in embryos (Parlak, 2018). New-generation pyrethroid derivatives, including etofenprox, were developed to reduce the toxic effects on fish (Matsuo, 2019). These compounds are specifically recommended for use in aquatic environments such as rice fields and for mosquito control purposes. While some ecotoxicological studies suggest that etofenprox is less toxic to fish (Matsuo, 2019), its effects on other aquatic organisms have been inadequately studied. Recent research has revealed acute toxic effects of etofenprox on algae, fish, and aquatic invertebrates (Sancho et al., 2018; Yaméogo et al., 2001; Zhang et al., 2010). There is a paucity of research on the sex-related endocrine-disrupting effects of these chemicals in non-model organisms, and most of the studied concentrations were higher than those found in the environment (Bovee et al., 2024; Eni et al., 2019; Souza et al., 2014). To our knowledge, no study has yet been published on the effects of etofenprox on amphibians at ecologically relevant concentrations, and neither deltamethrin nor etofenprox has been studied for their potential impact on amphibian sexual development. Given that mosquito control is specifically conducted in habitats and during periods where amphibian larvae develop in sensitive stages of sexual determination (Brühl et al., 2011), it would be especially relevant to investigate whether deltamethrin and etofenprox can cause sex reversal or sex-specific mortality in amphibians. This class of vertebrates is of particular concern for biodiversity conservation, with approximately half of their species being threatened with extinction (IUCN, 2024).

Here, we present an experimental study of the effects of deltamethrin and etofenprox applied at environmentally relevant concentrations during larval development on phenotypic sex ratio, sex reversal, and sex-specific mortality in agile frogs (*Rana dalmatina).* This anuran species has a male heterogametic sex determination system (XX/XY sex chromosomes), and a molecular marker set is available for diagnosing its genotypic sex (Nemesházi et al., 2020). Approximately 20% of phenotypic males are genotypically females in natural populations of agile frogs (Nemesházi et al., 2020), and female-to-male sex reversal can be induced by exposure to environmental stress during larval development (Mikó et al., 2021; Ujszegi et al., 2022). In the present study, we exposed tadpoles to one of the two pyrethroids in mesocosms during their most sensitive period of sex determination, mimicking the application of mosquito control, to test whether the treatments would trigger sex reversal or affect mortality in a sex-specific manner.

## MATERIALS AND METHODS

### EXPERIMENTAL PROTOCOL

All experimental procedures were approved by the Ethical Commission of the Plant Protection Institute, and permissions were issued by the Government Agency of Pest County (PE-06/KTF/00754-8/2022, PE-06/KTF/00754-9/2022, PE-06/KTF/00754-10/2022, PE/EA/295-7/2018). On April 29 in 2022, we collected ca. 150 agile frog eggs from each of seven freshly laid egg masses from a pond near Budapest, Hungary (Nagykovácsi Békás-tó 47°34’34.7“N, 18°52’08.1“E). Eggs were transported to the Experimental Station of the Plant Protection Institute (HUN-REN Centre for Agricultural Research) in Julianna-major, Budapest (47°32ʹ52ʺN, 18°56ʹ05ʺE). Until reaching developmental stage 25 (Gosner, 1960) on May 5, we reared each sibling group at a constant temperature of 18.5 ± 0.4 °C and a 14.5-9.5 hours light-dark cycle (corresponding to the natural photoperiod), housed separately in 5 L containers (24 × 16 × 13 cm) filled with 0.5 L reconstituted soft water (RSW; 48 mg NaHCO_3_, 30 mg CaSO_4_ × 2 H_2_O, 61 mg MgSO_4_ × 7 H_2_O, 2 mg KCl added to 1 L reverse-osmosis filtered, UV-sterilized, aerated tap water). When the hatchlings reached the free swimming state we placed the tadpoles into outdoor mesocosms. We set up mesocosms 6 weeks before the addition of tadpoles by filling 50 plastic tubs (42 × 72 × 30 cm) each with 65 L of tap water and leaving 5 days for chlorine to evaporate. Then we added 40 g dried beech (*Fagus sylvatica*) leaves into each mesocosm to provide nutrients and refuges for tadpoles. To enhance algal growth and establish a self-sustaining ecosystem, we inoculated each mesocosm two times (3 and 5 weeks before adding the tadpoles) with 1 L pond water containing phytoplankton and zooplankton. We covered mesocosms with mosquito screen lids to prevent colonization by insects. The 50 mesocosms were arranged in a randomized block design where no treatment was repeated within the same bloc. We distributed 700 tadpoles evenly across 5 treatment groups, resulting in 10 mesocosms in each treatment group. Each mesocosm contained 14 tadpoles, 2 members of each sibling group. Four treatment groups each received either deltamethrin or etofenprox at a lower concentration of 0.03 μg/L or a higher concentration of 0.3 μg/L (Sigma-Aldrich; catalogue numbers 34094 & 45423). We chose these concentrations because 0.03 μg/L falls in the middle range of pyrethroid concentrations measured in surface waters (Tang et al., 2018) and 0.3 μg/L falls between the two lowest concentrations of deltamethrin for which sex-disrupting effects were reported in fish (Eni et al., 2019). As the pyrethroids were added to the water dissolved in a small amount of ethanol, the fifth group (solvent control) received the same amount of ethanol in their water (0.3 μl/l 96% ethanol). This ethanol concentration is much lower than those observed to harm anuran embryos or tadpoles (Fainsod and Kot-Leibovich, 2017; Peng et al., 2005; Taylor and Brundage, 2013). The chemicals were added into the mesocosms three times on days 1, 15, and 35 of the experiment (i.e. after placing the tadpoles into the mesocosms) to simulate chemical mosquito control which is typically applied at two-week intervals in our country during the late spring and summer months.

Individuals that reached developmental stage 42 (i.e. the emergence of forelimbs, start of metamorphosis) were transferred into a 45 L (56 × 39 × 28 cm) plastic box (one box for each treatment group) covered with a perforated lid and placed into a shady outdoor area. Boxes contained shallow aged tap water and were slightly tilted to provide both a water-covered as well as a dry area. Each animal spent roughly a week under these conditions until it fully resorbed its tail (developmental stage 46, completion of metamorphosis); during this time we monitored their survival daily and recorded all mortality events. When a dead individual was found, we preserved it in 96 % ethanol for genotypic sexing (note that phenotypic sexing is not reliable at this early life stage). It was not possible to track individual survival during larval development, because dead tadpoles decompose quickly in the aquatic environment, and finding each corpse in time would have required very frequent thorough examination of every mesocosm, causing excessive disturbance to the tadpoles potentially resulting in artificially high mortality. However, only 4% of tadpoles died (i.e., could not be recovered from the mesocosms; N=27), which resulted in 661 individuals that started metamorphosis. In addition, N=12 tadpoles did not start metamorphosis by July 8 and had to be euthanized as described below.

Out of the animals surviving to the end of metamorphosis, we chose 60 individuals from each of the five treatment groups; the rest of the froglets were returned to their pond of origin. To avoid the selection of individuals who had exhibited extreme early metamorphosis, we released the first 10 froglets from each treatment group and kept the next 60. The retained froglets were moved into 2-L individual rearing containers in the laboratory, with wet paper towels as a substrate and a piece of egg carton as a shelter. The temperature in the lab was set to 21 ± 0.5 °C and the photoperiod was 8-16 dark-light hours. Froglets were fed two times a week *ad libitum* with small (2–3 mm) crickets (*Acheta domesticus*), sprinkled with a 3:1 mixture of Reptiland 76280 (Trixie Heimtierbedarf GmbH & Co. KG, Tarp, Germany) and Promotor 43 (Laboratorios Calier S.A., Barcelona, Spain) growth supplement. There was only one spontaneous mortality event among the froglets; we replaced this individual with another from the same treatment group to keep the sample size at N=300.

We dissected the animals 6–8 weeks after metamorphosis (15-19 August) to identify phenotypic sex (Mikó et al., 2021). At this age, the gonads are well differentiated in this species (Bernabò et al., 2011; Ogielska and Kotusz, 2004). When a froglet reached the desired age, we euthanized it using a water bath containing 6.6 g/L MS-222 (Sigma-Aldrich, catalogue number: E10521) buffered to neutral pH with the same amount of NaH_2_PO_4_. We examined the gonads under an APOMIC SHD200 digital microscope. We categorized phenotypic sex as male (testes), female (ovaries), or uncertain (abnormally looking gonads). We took a tissue sample (hind feet) from each froglet with flame-sterilized equipment and stored it in 96 % ethanol for genotypic sexing. We fixed the rest of the body in 13 ml neutral-buffered 4 % formalin (Sigma-Aldrich 1.00496), from which we later removed the gonads for histological analysis to clarify phenotypic sex where it was uncertain by gonad morphology.

### SEX IDENTIFICATION

Following the manufacturer’s protocol, we extracted DNA using E.Z.N.A. Tissue DNA Kit (Omega Bio-tek), except that the digestion time was at least 3 hours. For genotypic sexing, we used the method of (Nemesházi et al., 2020). Briefly, we used HRM (high resolution melting) analysis to test all samples for sex marker Rds3 (≥ 95% sex linkage; primers: Rds3-HRM-F and Rds3-HRM-R). The total HRM reaction volume was 15 µL, containing 3 µL “5x HOT FirePol EvaGreenPCR Mix Plus (ROX)” (Solis BioDyne), 1 µL forward and 1 µL reverse primer (10 µM each), and 100 ng genomic DNA. in MQ water to reach the final volume. Reactions were performed in a Roche LightCycler® 96 qPCR Instrument and the results were analyzed with the LightCycler 96 v.1.1.0.1320 software (Roche Diagnostics International LTD). For froglets dissected at the end of the study, we accepted an individual to be concordant male or concordant female if its Rds3 genotype was in accordance with its phenotypic sex. Individuals that appeared to be sex-reversed based on Rds3 genotyping were tested for another marker, Rds1 (≥89% sex linkage; primers: Rds1-F, Rds1-R, and Rds1-Y-R) using conventional PCR and were accepted to be sex-reversed only if both markers confirmed sex reversal. To study sex-specific mortality, we used the entire bodies of animals that died during metamorphosis and determined their genotypic sex using the Rds3 and Rds1 sex markers. If the results of these markers were consistent we accepted it as the genotypic sex of the animal. In the present study, there were no mismatches between the two markers.

### STATISTICAL ANALYSES

All statistical analyses were conducted in ‘R’ (version 4.2.3., R Core Team, 2023). We tested the effects of treatments on sex-specific mortality, sex-reversal rate, and phenotypic sex ratio by generalized linear models (GLM). We used diagnostic residual plots to ascertain that the data fit the statistical assumptions of each analysis (‘*simulateResiduals*’ function of the ‘DHARMa’ package (Badolo et al., 2018).

For the analysis of sex-specific mortality, we entered survival as the dependent variable (i.e. whether the individual survived metamorphosis or died) and the interaction of genotypic sex and treatment as fixed effects. To analyze the frequency of sex reversal we used Firth’s bias-reduced logistic regression (‘*brglmFit*’ function of the ‘brglm2’ package) (Kosmidis and Firth, 2021) because this yields less biased estimates when separation is present in the data (i.e. the number of sex-reversed individuals was zero in two treatment groups). We restricted this analysis to genotypically female individuals (XX genotypes), and the dependent variable was phenotypic sex (i.e. whether the individual was female-to-male sex-reversed or not). For the analysis of phenotypic sex ratio, we entered phenotypic sex as the dependent variable. We used the ‘*emmeans*’ function of the ‘emmeans’ package (Pike, 2011) to calculate odds ratios with standard errors from each model to express the differences of each pyrethroid treatment group from the control group (for phenotypic sex ratio and sex-reversal rate) and between males and females in each group (for mortality). For these pairwise comparisons, we present the *P*-values uncorrected as well as after adjusting with the false discovery rate (FDR) method (Pike, 2011). We report mean estimates with ± standard errors (SE).

## RESULTS

Out of the 113 individuals that died during metamorphosis and could be genotypically sexed, 72 were females and 41 were males. Out of the 543 individuals that survived metamorphosis, we chose 300 for further study; among the latter, there were 157 genotypic females and 143 genotypic males. Thus, genotypic sex ratio was significantly more female-biased among the animals that died (63.7%) than among the survivors (52.3%, Fisher’s exact test: *P* = 0.045). The GLM model showed that this sex-biased mortality was due to the low-concentration etofenprox treatment group, in which survival was significantly lower for females than for males (Table 1, Fig. 1). The latter difference in mortality was relatively large (i.e. the proportion of individuals that died was 0.39 ± 0.08 in females and 0.14 ± 0.05 in males; Fig. 1), although it became marginally non-significant after correcting the *P*-values for multiple comparisons (Table 1). There was no sex-biased mortality in the other treatment groups (Table 1, Fig. 1).

**Table (1).** Differences in mortality between sexes, expressed as odds ratios with standard errors (SE), in each treatment group (DTM_low_: deltamethrin 0.03 μg/L, DTM_high_: deltamethrin 0.3 μg/L, ETF_low_: etofenprox 0.03 μg/L, ETF_high_: etofenprox 0.3 μg/L, Control: 0.3 μl/L 96 % ethanol).

| Contrast | Treatment | Odds ratio ± SE | N | z | P |
| --- | --- | --- | --- | --- | --- |
| Female / Male | Control | 1.30 ± 0.58 | 94 | 0.57 | 0.565 |
| Female / Male | DTM<sub>low</sub> | 1.05 ± 0.57 | 77 | 0.09 | 0.921 |
| Female / Male | DTM<sub>high</sub> | 1.58 ± 0.79 | 83 | 0.89 | 0.369 |
| Female / Male | ETF<sub>low</sub> | 3.85 ± 2.11 | 82 | 2.45 | 0.014* |
| Female / Male | ETF<sub>high</sub> | 1.12 ± 0.61 | 77 | 0.21 | 0.830 |

\**P* = 0.070 after FDR correction for 5 comparisons.

**Fig.1.**
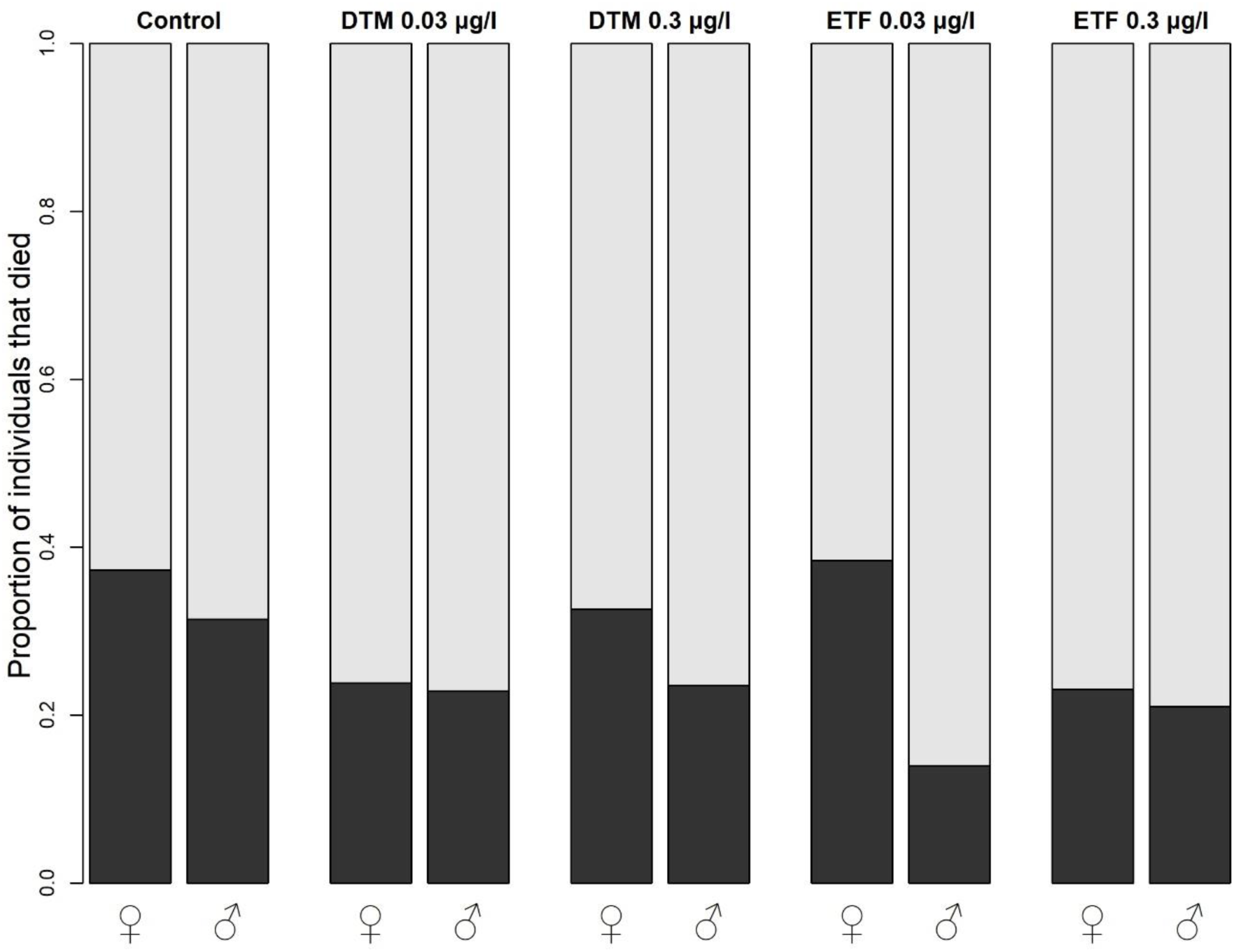
Mortality rate during metamorphosis by sex in different treatment groups (deltamethrin: DTM, etofenprox: ETF).

Among froglets dissected at the end of the study, we found a total of 9 female-to-male sex-reversed individuals: two in the control group, four in the low-concentration deltamethrin treatment, and three in the high-concentration etofenprox treatment (Fig. 2). Thus, the incidence of sex reversal was statistically independent of treatment (Table 2). Additionally, one genotypically female individual in the high-concentration deltamethrin treatment group had intersex gonads, i.e. ovotestes (Fig. 3).

**Fig.2.**
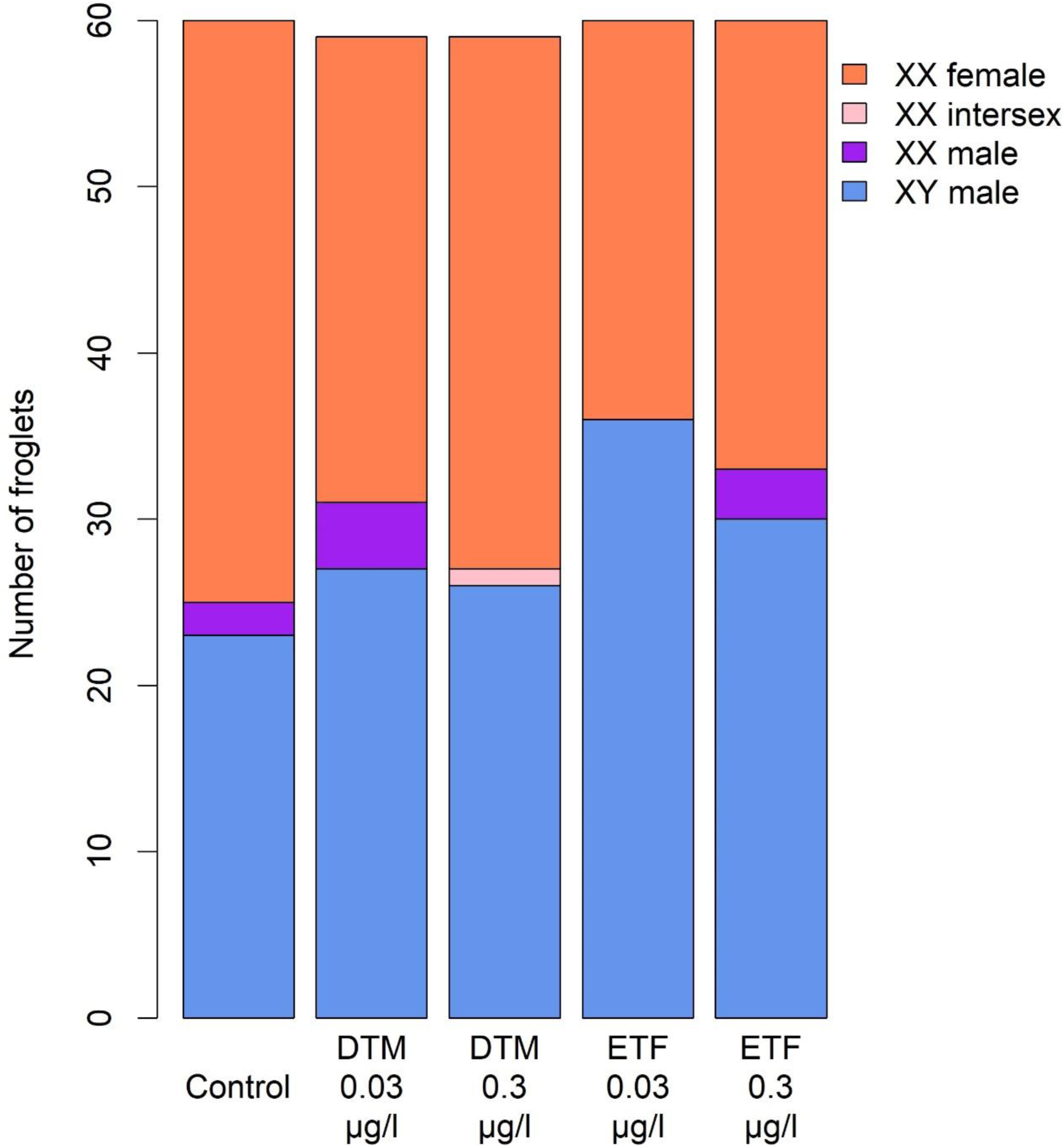
Distribution of combinations of genotypic and phenotypic sex among the treatment groups (deltamethrin: DTM, etofenprox: ETF). XX females and XY males, respectively, represent sex-concordant individuals, whereas XX males are female-to-male sex-reversed individuals, and XX intersex refers to a genotypically female individual with ovotestes.

**Table (2).** Effects of the treatments on sex-reversal rate, expressed as the odds ratio (with standard error, SE) of becoming phenotypic male in genotypic females. (Treatment groups: DTM_low_: deltamethrin 0.03 μg/L, DTM_high_: deltamethrin 0.3 μg/L, ETF_low_: etofenprox 0.03 μg/L, ETF_high_: etofenprox 0.3 μg/L, Control: 0.3 μl/L 96 % ethanol).

| Contrast | Odds ratio ± SE | z | P |
| --- | --- | --- | --- |
| DTM<sub>low</sub> / Control | 2.24 ± 1.88 | 0.96 | 0.336 |
| DTM<sub>high</sub> / Control | 0.21 ± 0.34 | -0.95 | 0.339 |
| ETF<sub>low</sub> / Control | 0.29 ± 0.46 | -0.77 | 0.439 |
| ETF<sub>high</sub> / Control | 1.80 ± 1.58 | 0.67 | 0.500 |

Sample sizes in each group: Control: 37, DTM<sub>low</sub>: 32, DTM<sub>high</sub>: 32, ETF<sub>low</sub>: 24, ETF<sub>high</sub>: 30

**Fig.3.**
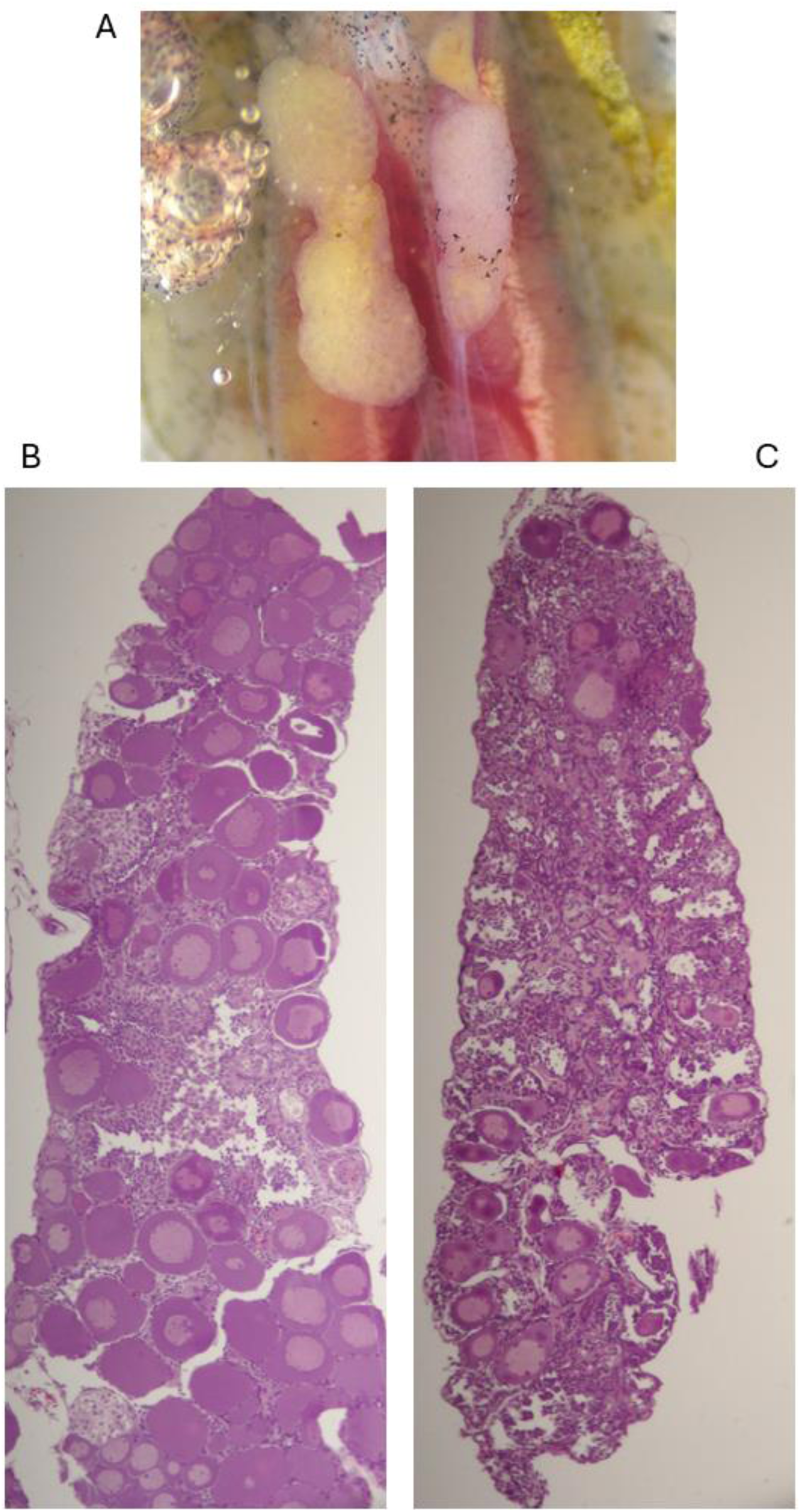
Photographs of the intersex gonads of an agile frog juvenile from the high-concentration deltamethrin treatment group: A) both gonads at dissection, B) histological sections of the right (B) and left (C) gonad.

Across all froglets, there were no significant differences in the phenotypic sex ratio between the pyrethroid treatment groups and the control group, except for the low-concentration etofenprox treatment group (Table 3, Fig. 2), where the proportion of males increased significantly relative to the control group’s sex ratio (Table 3). This shift in sex ratio was relatively large (i.e., from 0.42 17 ± 0.063 to 0.60 ± 0.06; Fig. 2), although it became statistically non-significant after correcting the *P*-values for multiple comparisons (Table 3).

**Table (3).** Effects of the treatments on phenotypic sex ratio, expressed as the odds ratio (with standard error, SE) of being phenotypic male, in all froglets. (Treatment groups: DTM_low_: deltamethrin 0.03 μg/L, DTM_high_: deltamethrin 0.3 μg/L, ETF_low_: etofenprox 0.03 μg/L, ETF_high_: etofenprox 0.3 μg/L, Control: 0.3 μl/L 96 % ethanol).

| Contrast | Odds ratio ± SE | z | P |
| --- | --- | --- | --- |
| DTM<sub>low</sub> / Control | 1.55 ± 0.57 | 1.18 | 0.235 |
| DTM<sub>high</sub> / Control | 1.14 ± 0.42 | 0.34 | 0.729 |
| ETF<sub>low</sub> / Control | 2.10 ± 0.78 | 1.99 | 0.045* |
| ETF<sub>high</sub> / Control | 1.71 ± 0.63 | 1.45 | 0.145 |

\**P* = 0.183 after FDR correction for four comparisons.
Sample sizes in each group: Control: 60, DTM<sub>low</sub>: 59, DTM<sub>high</sub>: 58, ETF<sub>low</sub>: 60, ETF<sub>high</sub>: 60

## DISCUSSION

In this study, we investigated the effects of environmentally relevant concentrations of deltamethrin and etofenprox on sex-dependent mortality, sex-reversal rate, and phenotypic sex ratio in agile frogs. We found that the lower concentration of etofenprox caused female-biased mortality and therefore shifted the phenotypic sex ratio towards males. We also identified female-to-male sex reversal in some individuals; however, this phenomenon was independent of the treatments. Neither the higher etofenprox concentration nor the deltamethrin treatments had significant effects on the measured variables. We discuss each of these findings in detail below.

Pyrethroids, including etofenprox, are known to have various biological effects that could explain the observed sex-specific mortality. One potential mechanism is the differential sensitivity of male and female individuals to oxidative stress. Studies on various taxa have shown that pyrethroids can induce oxidative stress, leading to cell damage and increased mortality (Ağırbaşlı et al., 2020; Nasia et al., 2018; Radovanović et al., 2017; Zhang et al., 2021). Oxidative stress can affect males and females differently due to sex differences in metabolic rates and antioxidant defences; females are usually less resistant to oxidative stress than males across vertebrates (Costantini, 2018). Another potential explanation is the interaction of pyrethroids with the endocrine system, which includes inhibiting steroid biosynthesis and modulating sex-hormone receptors (Wang et al., 2020). As steroid hormones are critical for regulating metamorphosis (Sachs and Buchholz, 2019), disruption of this process might have been responsible for the sex-dependent mortality observed during the metamorphic climax. The fact that this effect occurred only at the lower but not the higher concentration of etofenprox may be due to a non-linear dose-response relationship. The observation of non-monotonic dose-response effects in response to EDCs highlights that low doses can, in certain instances, have more significant effects than higher doses (Zoeller and Vandenberg, 2015). In our study, the female-biased mortality in the low-concentration etofenprox group might suggest a hormetic effect where low levels of a stressor cause more harm than higher levels. This non-linear response could be due to low-dose stimulation of oxidative stress pathways or endocrine disruption mechanisms that do not operate at higher concentrations. Although both the female-biased mortality and male-biased sex ratio in our low-concentration etofenprox treatment became statistically non-significant after adjusting the *P*-values for multiple testing, it must be borne in mind that all our response variables were proportions. The powerful analysis of proportions requires large sample sizes, making the results sensitive to penalizing by the number of comparisons. For example, had we tested only etofenprox (or both etofenprox and deltamethrin but only at the lower concentration) in our experiment, the same *P*-values would have remained statistically significant after FDR correction. Thus, the effects we found are non-negligible, especially given that the ecotoxicological effects of etofenprox are relatively poorly studied. However, more research is needed on etofenprox particularly on aquatic organisms, paying attention to potentially non-monotonic dose-response relationships and sex-dependent outcomes.

On the other hand, we cannot rule out that the effects we observed in the low-concentration etofenprox treatment were due to some random noise. For example, there was a heat wave between the end of June and the start of July, by which time most of the frogs had completed metamorphosis, but there were slightly more metamorphosing animals still in the outdoor container of the low-concentration etofenprox group than in the other treatment groups. Agile frogs are sensitive to heat stress, and although heat tolerance is not sex-dependent in the tadpole stage of the species (Bókony et al., 2024), sex difference in heat-related mortality might exist during the metamorphic climax, which is a critical time when mortality increases even under otherwise benign conditions (Bókony et al., 2020; Mikó et al., 2021). Therefore, the heat wave during the metamorphic climax could partly explain the female-biased mortality in the low-concentration etofenprox treatment. Thus, our etofenprox results might be attributable (at least in part) to chance events. This possibility is congruent with the rest of our findings, i.e. that neither the higher concentration of etofenprox nor any concentration of deltamethrin had any effect in our study. Altogether, this suggests that the ecologically relevant concentrations that we tested may not be harmful for agile frog tadpoles and metamorphs, which stands in contrast to earlier findings with higher concentrations on other taxa (Cengiz and Unlu, 2006; Eni et al., 2019; Macagnan et al., 2017; Vanzetto et al., 2019). This highlights that it is important to minimize environmental pollution by pyrethroids in order to prevent harmful effects.

Pyrethroids have been reported to act as androgen receptor antagonists and estrogen receptor modulators, disrupting normal hormonal signalling pathways essential for sexual development (Wang et al., 2020). For example, zebrafish (*Danio rerio*) embryos treated with 10 nM (5 μg/L) deltamethrin became male-biased, while lower concentrations produced slightly female-biased sex ratios (Bovee et al., 2024). In African sharptooth catfish (*Clarias gariepinus*), juveniles chronically exposed to 0.22–1.76 μg/L deltamethrin had ovotestes and increased estradiol and reduced testosterone concentrations in blood plasma (Eni et al., 2019). However, in our study, larval exposure to deltamethrin or etofenprox did not alter gonadal development in agile frogs, as these treatments did not increase the rate of sex reversal relative to the very low rates of sex reversal observed in the control group, which corresponds to the rare incidence of spontaneous sex reversal reported from this species (Bókony et al., 2024, 2021; Mikó et al., 2021; Ujszegi et al., 2022). Nevertheless, the presence of sex-reversed and intersex individuals in our pyrethroid treatments, and a potentially sex-reversing effect of beta-cypermethrin observed in *Trichogramma pretiosum* (Souza et al., 2014) tentatively suggest that pyrethroid exposure might disrupt sexual development under certain circumstances. Since, to our knowledge, no other studies examined this question, further investigations are needed to test this idea in various taxa such as amphibians, whose sex determination and sex development are sensitive to environmental stressors including chemical pollutants (reviewed by Nemesházi and Bókony, 2022).

Finally, our study highlights an important lesson for ecotoxicology in general. Although the phenotypic sex ratio of exposed young is an important endpoint in ecotoxicological experiments, by itself it cannot demonstrate an endocrine-disrupting effect on gonadal development. Had we studied only the phenotypic sex ratio here, we might have mistakenly concluded that the lower concentration of etofenprox may cause sex reversal in agile frogs. By analyzing genotypic sex in the surviving and perished animals, we could reveal that there was actually zero sex reversal in this treatment and, instead, sex-biased mortality was responsible for the shift in phenotypic sex ratio. Teasing apart these underlying mechanisms has long been hindered by the lack of genotypic sexing methods for the majority of ectothermic vertebrate species; this limitation forced most earlier ecotoxicological studies to make inferences solely based on phenotypic sex ratios (Nemesházi and Bókony, 2022). However, with the increasing availability of genomics techniques, genotypic sex markers are being published on more and more species, which will help raise the ecotoxicological research of “sex disrupting” effects to a higher level. A better understanding of these effects is crucial for the assessment of the long-term impacts of environmental contaminants on biodiversity and for the development of effective conservation strategies.

## AUTHOR CONTRIBUTIONS STATEMENT

Emese Balogh: Methodology, Investigation, Data curation, Formal analysis, Writing - original draft, Writing - review & editing, Visualization, Funding acquisition. Szabolcs Hócza: Investigation, Data curation, Writing - review & editing. Nikolett Ujhegyi: Investigation, Data curation, Writing - review & editing. Andrea Kásler: Investigation, Writing - review & editing. Dóra Holly: Investigation, Writing - review & editing. Dávid Herczeg: Investigation, Writing - review & editing. János Ujszegi: Investigation, Writing - review & editing. Zoltán Gál: Investigation, Writing - review & editing. Orsolya I. Hoffmann: Investigation, Funding acquisition, Writing - review & editing. Veronika Bókony: Conceptualization, Methodology, Investigation, Formal analysis, Writing - review & editing, Supervision, Funding acquisition, Data curation and Visualization. Zsanett Mikó: Conceptualization, Methodology, Investigation, Data curation, Writing - review & editing, Supervision, Funding acquisition. All authors contributed critically to the drafts and gave final approval for publication.

## ACKNOWLEDGEMENTS

We are thankful to Csenge Kalina, Anna Kraxner and Márk Szederkényi for their help during the experiment and data archiving. We thank Botond Pertics and Ferenc Samu at the NÖVI Department of Zoology for providing us with their RT-PCR machine and other lab facilities. The study was funded by the National Research, Development and Innovation Office of Hungary (NKFIH, grants 135016 to V.B. and 134241 to Z.M.). The authors were supported by the strategic research fund of the University of Veterinary Medicine Budapest (Grant No. SRF-001 to E.B. and V.B.), the New National Excellence Program (ÚNKP-21-3, ÚNKP-22-3 to A.K. and ÚNKP-23-4 to A.K. and J.U.) and the Egyetemi Kutatási Ösztöndíj Program of the Ministry for Innovation and Technology from the source of the National Research, Development and Innovation Fund (EKÖP-24-3 to E.B., EKÖP-24-4 to Z.M., N.U. and J.U.). H.D. gained support from the János Bolyai Research Scholarship of the Hungarian Academy of Sciences (MTA, BO/00549/24/8). This project received funding from the HUN-REN Hungarian Research Network.

## DATA AVAILABILITY

Data will be made available on request.

